# *In vitro* modeling of tumor spheroid interactions to perfused blood vessels

**DOI:** 10.1101/2020.08.03.234633

**Authors:** Tae Joon Kwak, Esak Lee

## Abstract

Tumor angiogenesis, the formation of new blood vessels from existing blood vessels in and around the tumor stroma, is orchestrated by multiple biological factors in the tumor microenvironment, including tumor stromal cells, extracellular matrices, and secreted growth factors. Here, we present the processes to define and optimize the biological conditions for robust interactions between tumor spheroids and engineered blood vessels in a microfluidic organ-on-a-chip device *in vitro*. Within the device, vascular lumen formation and vessel sprouting in human umbilical vein endothelial cell based engineered blood vessels are observed in a variety of extracellular matrices (collagen, Matrigel, and fibrin), containing metastatic breast tumor cells (MDA-MB-231), and tumor stromal cells (mesenchymal stem cells, and human lung fibroblasts) to show the crosstalk between the tumor spheroids and the perfused blood vasculatures *in vitro*.

## Introduction

Cancer is one of the leading causes of human death in modern society. Tumor angiogenesis, a process of new blood vessel formation from existing blood vessels in and around the tumor stroma has been appreciated as one of the major targets for treating cancer (Folkman, 1971). Tumor blood vessels are morphologically and functionally different from physiological blood vessels, which may affect tumor oxygenation, metabolism, and anti-cancer drug delivery and tumor drug responses (Lee, Pandey, & Popel, 2015). Besides, to better understand and treat tumor metastasis or tumor cell dissemination to distant organs, we need to look into interactions between the tumor mass and nearby blood vessels (Lee et al., 2015).

To model a tumor mass *in vitro*, tumor spheroids have been used (Fennema, Rivron, Rouwkema, van Blitterswijk, & de Boer, 2013). Tumor spheroids are clusters of tumor cells formed due to the tendency of adherent cells to aggregate when growing in the media suspension (Fennema et al., 2013). Multicellular tumor spheroids have been used to model human solid tumors because of their morphological and biological similarities to *in vivo* human tumors. Indeed, compared to two-dimensional (2D) tumor cell culture models, three-dimensional (3D) multicellular tumor spheroids could provide more realistic cell-cell, cell-extracellular matrix (ECM) interactions with multiple stromal cells, which are the essential characteristics of the heterogeneous tumor microenvironment (TME) (Chung, Ahn, Son, Kim, & Jeon, 2017; Hirschhaeuser et al., 2010; Ko et al., 2019; Nashimoto et al., 2020). In particular, spheroids models have been useful because they could display morphological and size changes in 3D when the cancer cells’ traits are transformed in the ECM under anti-cancer drug treatments (Antoni, Burckel, Josset, & Noel, 2015). The 3D tumor spheroids models are thus a relevant model for tumor cell growth, migration, and differentiation (Antoni et al., 2015).

*In vitro* modeling of cancer progression has been emerging (D.H.T. Nguyen et al., 2019; Hassell et al., 2017; Lee, Song, & Chen, 2016). Recently, examining the relationship between blood vessels and tumor spheroids in 3D *in vitro* devices has attracted attention because the models have offered more realistic environments (e.g., dimensionality, fluid flow (luminal/interstitial), physical deformation, and ECM stiffness) to simulate *in vivo* human body than conventional cell culture on 2D plastic. For instance, therapeutic efficacies of anti-cancer drugs were assessed in tumor spheroid models under controlled fluid flow in an engineered tumor vascular network (Nashimoto et al., 2020). In this study, the anti-cancer drugs under the flow did not show dose-dependency that has been observed under the static condition. In addition, the proliferation of tumor cells in the spheroids were more accelerated under the flow perfusion (Nashimoto et al., 2020). In another study, differentiation and migration of vascular endothelial cells were observed under pro-angiogenic growth factor treatments, and permeability of the angiogenic sprouts within the collagen matrix was measured (van Duinen et al., 2019). However, it is still challenging to reproduce an appropriate *in vitro* model system of perfused blood vessels and their crosstalk to solid tumors with various environments (e.g., ECM, stromal cells, growth factors).

In this work, we demonstrate the optimization process to define the biological determinants of successful interactions of tumor spheroids with perfused blood vessels in a microfluidic organ-on-a-chip device *in vitro* to mimic vascularized solid tumors *in vivo*. Within our device, vascular lumen formation and vessel sprouts of the engineered blood vasculatures are observed in a variety of ECM environments containing metastatic tumor cells and supporting stromal cells to show the crosstalk between the tumor spheroids and the blood vasculatures *in vitro*.

## Materials and methods

### Cell culture

Human umbilical vein endothelial cells (HUVECs) were purchased from Lonza and were cultured in EGM-2 media (Lonza, Switzerland). MDA-MB-231 breast cancer cells were purchased from ATCC and cultured in DMEM (low glucose) + 10% Fetal bovine serum (FBS) + 2 mM L-glutamine + 50 µg/ml Gentamycin. Human mesenchymal stem cells (MSCs, bone marrow-derived) and human lung fibroblasts (HLFs) were purchased from Lonza and cultured in DMEM (low glucose) + 10% FBS + 2 mM L-glutamine + 50 µg/ml Gentamycin. Endothelial cells were used at passages 3-8, and the stromal cells (MSCs and HLFs) were used at passages 3-6. To achieve labelled Green Fluorescent Protein MDA-MB-231 (GFP-MDA-MB-231) cells, the MDA-MB-231 cells were transduced with a lentiviral construct pCSCG-EGFP (Addgene). To achieve labelled mApple Red Fluorescent Protein HUVECs (mApple HUVECs), the HUVECs were transduced with a lentiviral construct pBAD-mApple (Addgene). All the cells were maintained in standard tissue culture incubators at 37°C, 95% humidity, and 5% CO_2_.

### Device fabrication

The Polydimethylsiloxane (PDMS) casting mold was fabricated by following microfabrication method of Polacheck et al (Polacheck, Kutys, Tefft, & Chen, 2019), and the PDMS microfluidic chip device was fabricated using conventional PDMS casting process (Kwak et al., 2016). Briefly, a 10:1 ratio mixture of PDMS polymer and cross-linker curing agent (SYLGARD™ 184 Silicone Elastomer Kit, The Dow Chemical Company, MI, USA) was degassed, cast into the PDMS casting mold, and left in an oven for at least two hours at 80°C. The polymerized PDMS device was punched for inlets and outlets of media reservoir and ECM ports region, and bonded to a glass coverslip with a plasma reacted ion etch (RIE) surface treatment (PE-25 Plasma Cleaner, Plasma Etch Inc., NV, USA). The volume of the ECM cavity in the center of the microfluidic chip device is 20 mm^3^, and the distance between the ECM cavity and the media reservoir is 4 mm. Overall design and dimension of the microfluidic chip device is illustrated in Figure 1.

**Figure 1.**
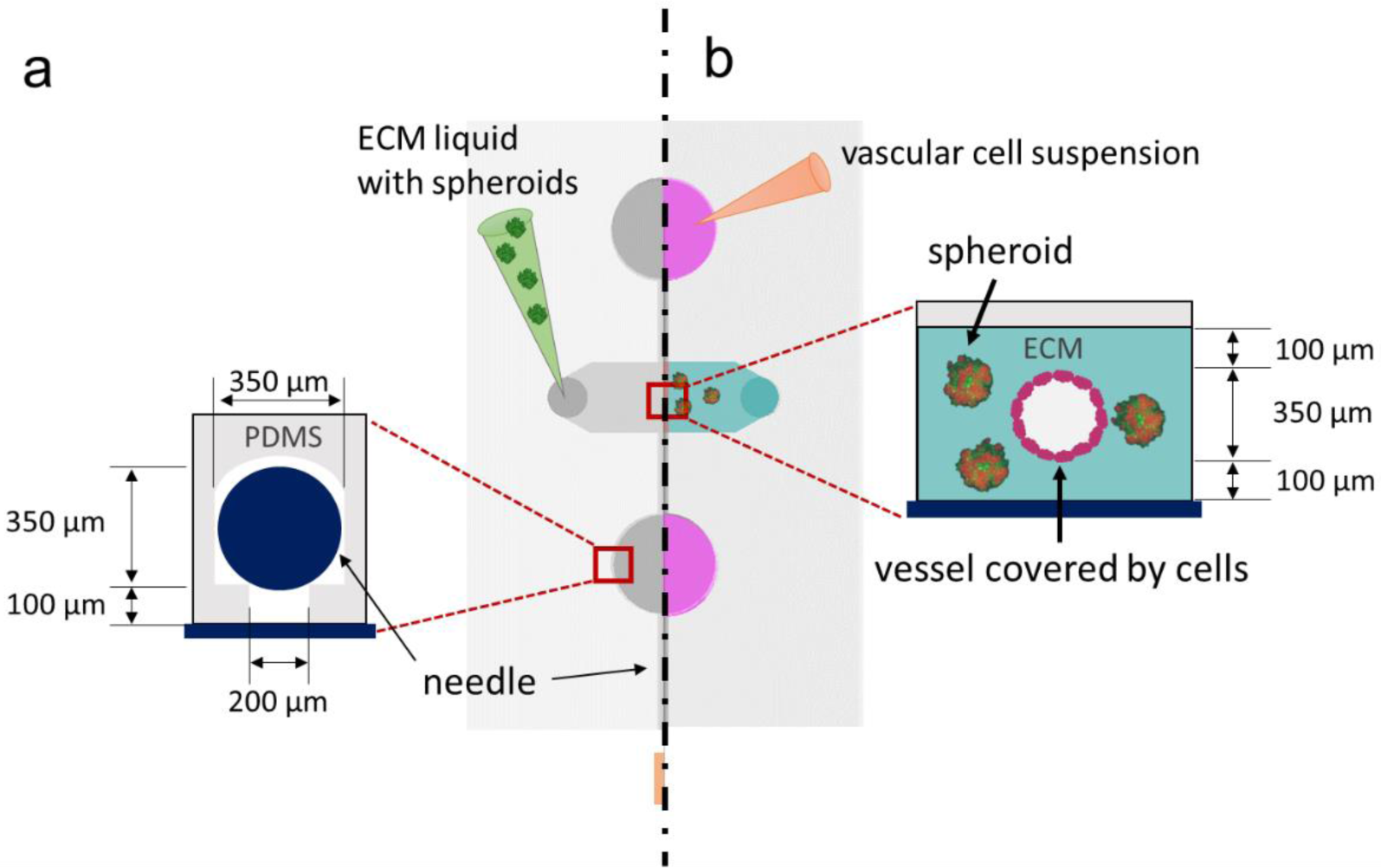
ECM casting with tumor-spheroids and endothelial cell seeding. (a) A casting needle is introduced into the channel in the microfluidic chip device, and the ECM hydrogel liquid is injected with tumor-spheroids and is polymerized as a gel around the needle. (b) The casting needle is removed after gelation to create an empty channel embedded in the ECM. Then, HUVECs are introduced into the channel via the media port and allowed to adhere to the ECM channel.

### Tumor spheroid preparation

MDA-MB-231 breast cancer cells (ATCC) under normal culture in DMEM were rinsed with phosphate-buffered saline (PBS) twice and detached by using trypsin. Trypsinized and detached MDA-MB-231 cells were diluted with excess culture media (DMEM) and centrifuged at 1,000 x g for 4 min. The supernatant was removed, and the MDA-MB-231 cell pellet was reconstituted with fresh DMEM to obtain 1.5 x 10^4^ cells/ml. The cell suspension was transferred into Ultra-Low Attachment (ULA) 96-well round-bottom plates (Sigma). Cell suspension of 200 µl was loaded in each well of the 96-well plate, loading 3,000 MDA-MB-231 cells per well. In the case that we co-culture stromal cells with the MDA-MB-231 tumor cells in the spheroids, we fixed the number of MDA-MB-231 tumor cells at 3,000 cells per well, and only the stromal cells were added with different ratios. For example, stromal cells cocultured with the tumor cells in the spheroids include (1) MSCs (MDA-MB-231:MSCs=1:1 or 1:3); (2) HLFs (MDA-MB-231:HLFs=1:1); (3) HUVECs (MDA-MB-231:HUVECs=1:2). The 96-well plates were incubated at 37°C, 5% CO_2_, 95% humidity and the spheroid formation was observed daily.

### ECM casting with tumor spheroid

The microfluidic chip device was plasma RIE treated and immersed in 0.1% Poly-L-Lysine PLL (#0413, ScienCell Research Laboratories, CA, USA) for 4 hours and rinsed with DI water three times. Subsequently, 1% glutaraldehyde (G6257, Millipore Sigma, MO, USA) was added to the ECM port and treated for 15 minutes, rinsed with DI water three times, and was soaked overnight in the DI water on an orbital shaker to fully remove any excess glutaraldehyde. The tumor spheroids (e.g., MDA-MB-231 and MSC, MDA-MB-231 and HLF, MDA-MB-231 and HUVEC, and only MDA-MB-231) were cultured and collected from 96-well plate as described above in the Tumor spheroid preparation. To mold cylindrical channel in the ECM, a 350 µm diameter of 1% bovine serum albumin (BSA) treated casting needle was inserted into the needle guide channel and ECM liquid materials i.e. rat tail collagen type I (354236, Corning, NY, USA), Matrigel^®^ (08-774-392, Corning, NY, USA), and fibrinogen (F8630, Millipore Sigma, MO, USA) plus thrombin (Sigma) with the tumor spheroids were mixed and polymerized in the cavity of the device around the casting needle, respectively as shown in the Figure 1a. For some experiments, we loaded human lung fibroblasts (HLFs) with density of 500,000 cells/ml in addition to the tumor spheroids in the ECM casting procedure to promote vascular sprouting from the vascular channel and tumor spheroids.

### Vascular cell seeding into devices

Once the ECM materials with tumor spheroids were solidified, the casting needle was removed to leave an empty cylindrical channel completely embedded in the ECM. Afterwards, 1.4 ml of HUVECs were seeded into the channel through the reservoir (cell seeding density at 1 million cells/ml) and allowed to form a monolayer cylindrical vascular channel as shown in the Figure 1b. To apply physiologically pertinent laminar shear stresses (>3 dyne/cm^2^), the cell-seeded microfluidic chip device was placed on a platform rocker with −30° to 30° angle at 3 rpm to generate gravity-driven luminal flow through the vascular channel at 37°C, 5% CO_2_, 95% humidity. The culture media (EGM-2) in the device was replenished daily.

### Immunoblot assays

Proteome Profiler Antibody Array Kits for human angiogenesis factors (R&D systems, MN, USA) were used for reverse western blot. We followed the manufacturer’s operation manual to investigate the relative levels of angiogenesis-related proteins in endothelial growth media-2 (EGM-2) and human lung fibroblast conditioned media (HLFCM).

### Quantification of spheroid growth pattern and sprouting/invasion length

The normalized area of each growth pattern was analyzed using ImageJ image analysis software (v1.52a, NIH, USA) software with fluorescence color channel splitting, thresholding, and particle analysis module (Kopanska et al., 2015). The average sprouting length was also quantified using ImageJ by manual determination of the furthest distance between the tumor spheroids and the tip of sprouting/invasion.

### Statistical analysis

Independent sample populations were compared using unpaired, two-sample t-test with a normal distribution assumption. *P < 0.05 was the threshold for statistical significance. All the error bars depict standard deviation.

## Results

### Migration of tumor spheroid mixtures

We first evaluated the growth pattern of the spheroid mixture of MDA-MB-231 metastatic breast cancer cells and mesenchymal stem cells (MSCs), given that MSCs are known to promote tumor progression in the tumor microenvironment (TME) (Atiya, Frisbie, Pressimone, & Coffman, 2020). Tumor spheroid mixtures groups were: (1) monoculture of MDA-MB-231 cells, (2) a 1: 1 ratio of MDA-MB-231 cells and MSCs coculture, and (3) a 1: 3 ratio of MDA-MB-231 cells and MSCs coculture as shown in Figure 2a. These tumor spheroids were cultured in DMEM containing Matrigel in 96 wall plates for 5 days. To assess the tumor growth pattern, the area changes in each spheroid were daily traced by normalizing the area by the initial area of the spheroid using the ImageJ software (Figure 2b). By comparing the growth patterns, the 1: 1 ratio mixture spheroid of MDA-MB-231 metastatic breast cancer cells and MSCs showed the fastest growth for 5 days among the groups.

**Figure 2.**
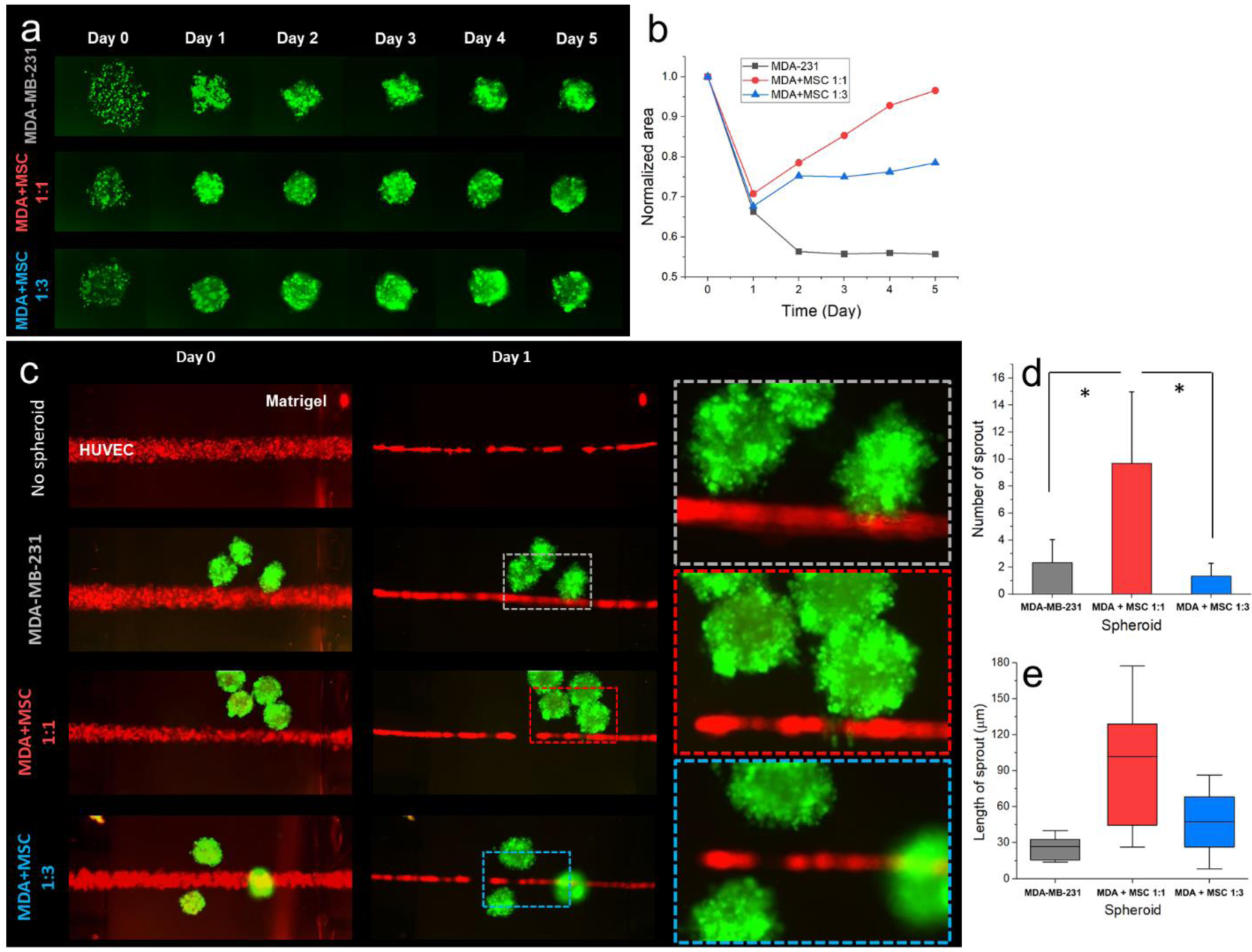
Growth and invasion of tumor spheroid mixtures. (a) Growth pattern of tumor spheroid mixtures (monoculture of MDA-MB-231 and coculture with MDA-MB-231 and MSCs) for 5 days. (b) Normalized growth area plot of the tumor spheroids for 5 days. (c) Tumor spheroids-HUVEC vasculature interaction in a microfluidic chip device. Enlarged images of each highlighted spheroids area are given the right side of (c). The number of sprout (n=3, respectively) (d) and the length of sprouts (e) of each spheroid were quantified under each condition. Green cells represent GFP-expressing MDA-MB-231; red cells represent mApple-expressing HUVECs.

Next, each of the tumor spheroids was introduced into the microfluidic chip device cast by Matrigel to investigate tumor spheroids-HUVEC channel interactions as shown in Figure 2c. Each tumor spheroid formed invasive sprouts to the HUVEC blood vessel, and the number and the length of the sprouts were quantified and plotted on Figure 2d and e. By comparing the sprout growth patterns, the 1: 1 ratio mixture spheroids of MDA-MB-231 metastatic breast cancer cells and MSCs showed the highest number of sprouts with the longest invadopodium among the three conditions. Interestingly, all the HUVEC vascular channels, with or without spheroids, were collapsed in the Matrigel ECM environment at Day 1 (Figure 2c). The collapsed vessels did not allow media perfusion through the vascular lumen. These data demonstrate that Matrigel negatively affects lumen formation in blood endothelial cells.

### Spheroids with MDA-MB-231 and MSC in different ECM

As observed in Figure 2c, the tumor spheroids comprising MDA-MB-231 metastatic breast cancer cells and MSCs at the 1:1 ratio showed the most aggressive growth and invasion pattern, while HUVEC vascular channels were collapsed in Matrigel ECM. Thus, we next explored different ECM conditions with the 1:1 mixture of MDA-MB-231 cells and MSCs tumor spheroids. As shown in the Figure 3, the microfluidic chip device embedded 5 different ECM conditions: (1) only collagen I, (2) only Matrigel, (3) 1:1 mixture of Matrigel and collagen I (denoted as M-gel and C-gen), (4) 1:2 mixture of Matrigel and collagen I, and (5) 2:1 mixture of Matrigel and collagen I, respectively. Each tumor spheroid formed invasive sprouts to the HUVEC blood vessel, and the number and the length of the sprouts were quantified and plotted on Figure 3b and c. In the collagen I dominant ECM microenvironments, such as the only collagen I and the 1:2 mixture of Matrigel and collagen I, the HUVEC vasculatures were not collapsed, enabling the media perfusion through the vascular lumens, however, the tumor invasion/sprouting behaviors in the tumor spheroids were less aggressive than those in the Matrigel dominant ECM conditions, such as the only Matrigel or the 2:1 mixture of Matrigel and collagen I. In the case of the only collagen I, none of the sprout to the HUVEC channel was found, and the average number of sprouts in the 1: 2 mixture of Matrigel and collagen I was 12.67 ± 6.13 which is slightly more than that in the cases of the 1: 1 and 2: 1 mixture of Matrigel and collagen I which were 12 ± 5.72 and 10.67 ± 4.03 respectively (Table 1 and Figure 3b); However, the average length of sprouts In the case of the 1: 2 mixture of Matrigel and collagen I was 24.75 ± 11.14 µm, which is definitely shorter than that in the cases of the 1: 1 and 2: 1 mixture of Matrigel and collagen I which were 40.48 ± 27.23 µm and 82.81 ± 38.44 µm, respectively (Table 2 and Figure 3c). By contrast, in the Matrigel dominant ECM conditions, the HUVEC vasculatures were all collapsed, although they have relatively more aggressive sprouts, as similar to the results in Figure 2. Consequently, we noticed that none of these ECM conditions tested simultaneously showed both tumor spheroids invasion/sprouts actively and the HUVEC vasculatures are successfully maintained (e.g., lumen formation, media perfusion).

**Table1.** Number of sprout.

| ECM | Collagen I | Matrigel | M-gel+C-gen<br>1:1 | M-gel+C-gen<br>1:2 | M-gel+C-gen<br>2:1 |
| --- | --- | --- | --- | --- | --- |
| Number of sprout | n/a | 21 ± 7.58 | 12 ± 5.72 | 12.67 ± 6.13 | 10.67 ± 4.03 |

**Table2.** Length of sprout.

| ECM | Collagen I | Matrigel | M-gel+C-gen<br>1:1 | M-gel+C-gen<br>1:2 | M-gel+C-gen<br>2:1 |
| --- | --- | --- | --- | --- | --- |
| Number of sprout | n/a | 29.63 ± 15.52 | 40.48 ± 27.23 | 24.75 ± 11.14 | 40.46 ± 18.78 |

**Figure 3.**
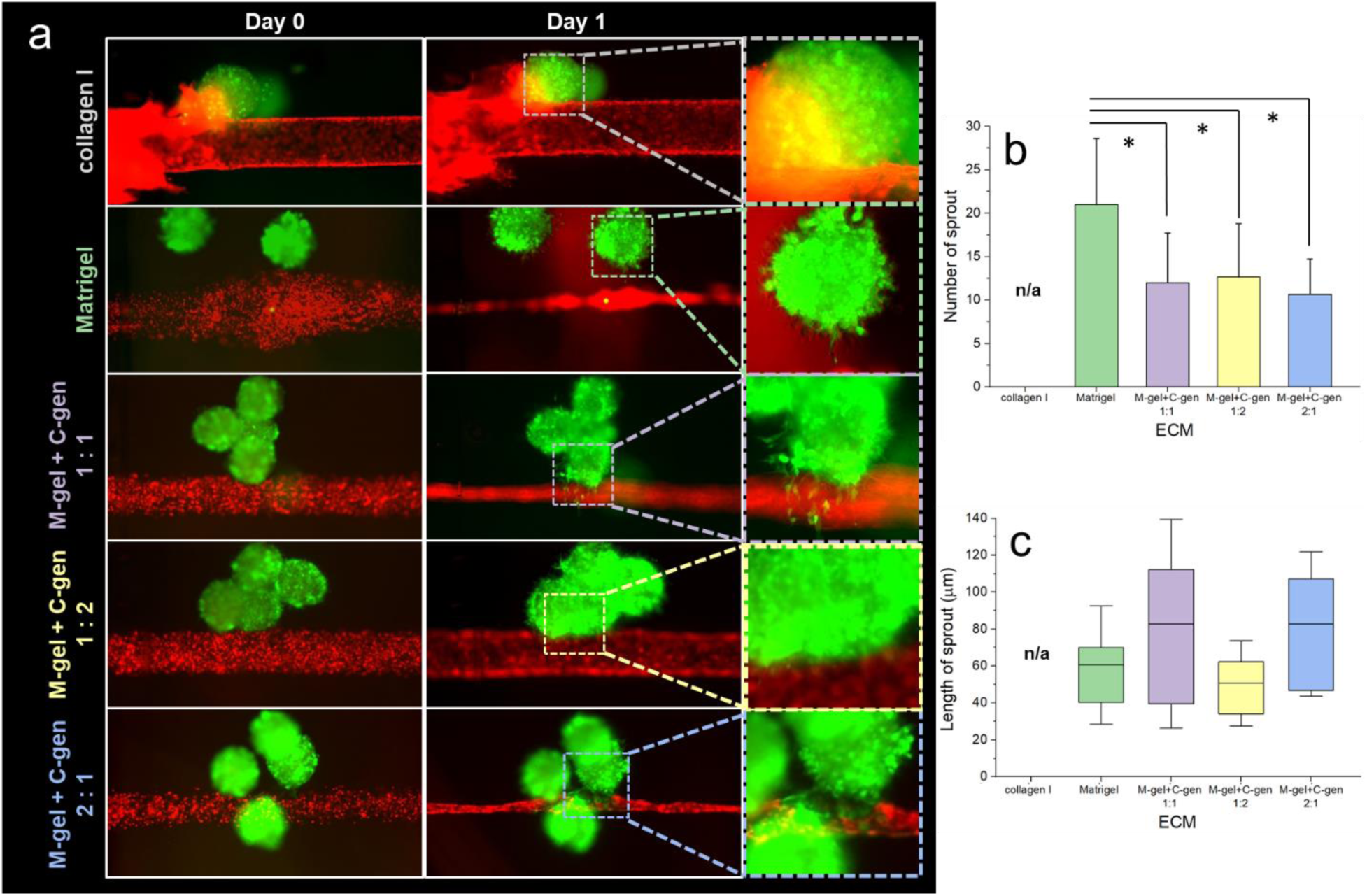
Spheroids with MDA-MB-231 and MSCs in different matrices. (a) The 1:1 mixture of MDA-MB-231 metastatic breast cancer cells and MSCs tumor spheroids were introduced into the microfluidic chip device embedded in different ECM microenvironment. Enlarged images of each highlighted spheroids area are given the right side. The number of sprout (b) and the length of sprouts (c) of each spheroid were quantified under each condition (n=3, respectively). Green cells represent GFP-expressing MDA-MB-231; red cells represent mApple-expressing HUVECs.

### Tumor spheroid-induced angiogenesis

Since we could not simultaneously demonstrate both the tumor spheroids sprout and the perfusable HUVEC vascular lumen in the collagen I and/or Matrigel ECM, as conducted in the above trials, we further explored the fibrin gel embedded ECM microenvironment in the microfluidic chip device. Since it has been reported that integrins in tumor cells and endothelial cells interact with fibrin, facilitating tumor invasion and angiogenesis and synergize pro-angiogenic growth factors, such as fibroblast growth factor 2 (FGF-2) and vascular endothelial growth factor (VEGF) (Laurens, Koolwijk, & De Maat, 2006; Lee et al., 2015). In addition, we considered additional stromal cell, human lung fibroblasts (HLFs) given the stromal cell support vascular sprouting and tumor invasion (Kim, Chung, & Jeon, 2016; Lee et al., 2015; Lee et al., 2016). We hypothesized that HLFs in fibrin could not only induce more active angiogenesis but also maintain vascular lumens. In this trial, tumor spheroids mixtures of MDA-MB-231 cells (denoted as MDA-231) + HLFs (Figure 4) were introduced for the interaction between the tumor spheroids and the HUVEC vasculatures. Particularly, for better mimicking of the realistic TME in the human body, we introduced HLFs as a growth factor supplier in the ECM bulk in addition to the tumor spheroids. Interestingly, the mixture of tumor spheroids with MDA-MB-231 cells and HLFs were actively sprouting while the HUVEC vasculature was not collapsed and showed angiogenic sprouting during 6 days of the experiment (Figure 4c-e). The average number and length of the sprouts in this trial were 42 ± 9.42 and 196.08 ± 59.64 µm, respectively. However, although both tumor spheroids and the vasculature were actively sprouted, they had not invaded each other.

**Figure 4.**
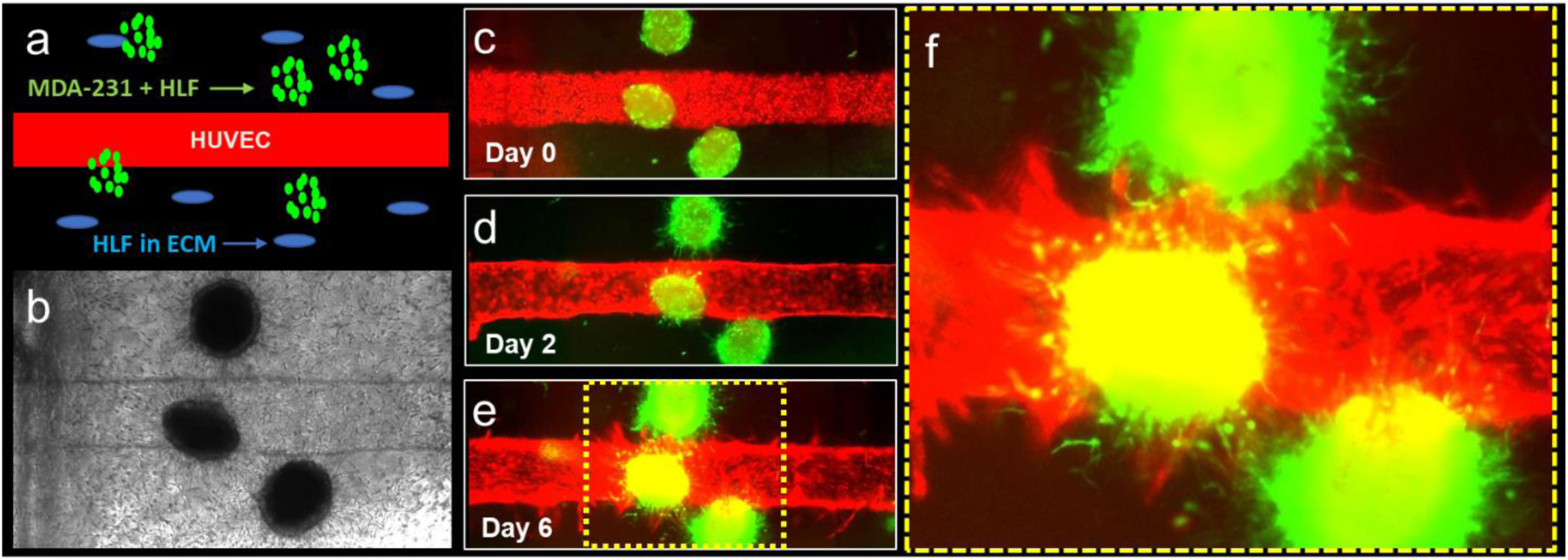
MDA-231 and HLF tumor spheroid-induced angiogenesis. (a) A schematic illustration of the experimental design. (b) A representative bright-field image of the HUVEC channel and tumor spheroids in the HLF laden ECM in the microfluidic chip device. (c-d) Confocal images of MDA-231 and HLF tumor spheroid-induced angiogenesis at day 0 (c), day 2 (d), and day 6 (e). At the same time tumor cells invade the ECM (f) An enlarged tumor spheroid-vasculature area highlighted in (e).

We further investigated the growth behavior of the tumor spheroids mixture of MDA-MB-231 cells + HUVECs in the HLFs laden fibrin ECM as displayed in Figure 5a, b. Since we found that the growth behavior of the tumor spheroids mixture of MDA-MB-231 cells + HUVECs is significantly active in the given ECM for 5 days (Figure 5c, d), we introduced the MDA-MB-231 cells + HUVECs tumor spheroids into the microfluidic chip device with a HUVEC channel to see the angiogenic interaction between the spheroids and the engineered blood vessel. Noteworthily, the mixture of tumor spheroids with MDA-MB-231 cells and HUVECs were actively sprouting out while the HUVEC vascular lumens were successfully maintained and showed angiogenesis for a total of 10 days of the experiment (Figure 5e-g) with the average number and length of the sprouts of 26.5 ± 3.5 and 273.59 ± 43.02 µm, respectively. Moreover, the sprouted tumor spheroids and HUVEC vascular channels have been invading each other since day 6 (Figure 5e-h), as confirmed by the enlarged confocal image (Figure 5i) with reversed pseudo colored one (Figure 5j).

**Figure 5.**
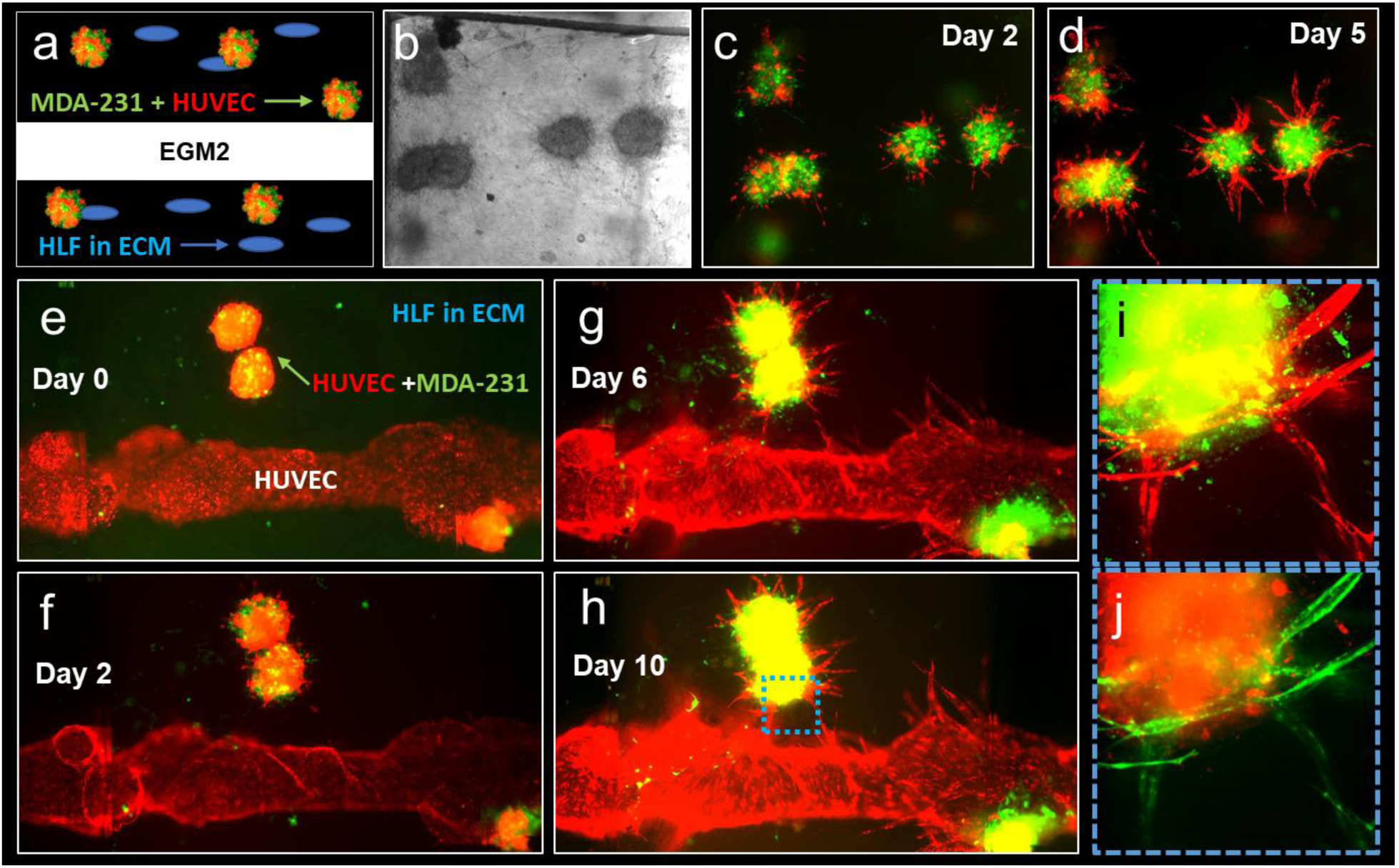
MDA-231 and HUVEC tumor spheroid-induced angiogenesis. (a-d) Growth of tumor spheroid mixture of MDA-231 and HUVEC. (a) A schematic illustration of the experimental design. (b) A representative bright-field experimental image of the tumor spheroids and HLFs laden in the ECM in the microfluidic chip device supplemented with EGM-2. (c-d) Representative confocal images of the spheroids of MDA-231 and HUVEC at day 2 (c), day 5 (d), respectively. (e-h) Representative confocal images of MDA-231 and HUVEC tumor spheroid-induced angiogenesis at day 0 (e), day 2 (f), day 6 (g), and day 10 (h), respectively. (e, f) Enlarged cross fluorescence confocal images of tumor spheroid-vasculature area highlighted in (h) as original fluorescence (i) and reversed pseudo-colored one (j).

Subsequently, we investigated how HLFs induced angiogenesis and maintained vascular lumens. The HLF conditioned media (HLFCM) was prepared as described previously (Lee et al., 2014) to examine the angiogenesis-related protein expression profiles contributed to each experiment. Briefly, when HLFs reached confluence in T175 tissue culture flasks, the normal growth media were replaced with 8 ml serum-free media. After 24 h incubation, the supernatant was centrifuged and filtered through 0.2 mm syringe filters (Corning). The resulting HLFCM were stored in aliquots at −80°C to avoid multiple freeze-thaws. The Figure 6 shows results of the membrane-based antibody array for the determination of the relative levels of angiogenesis-related proteins in EGM-2 (Figure 6a) and HLFCM (Figure 6b). Each specimen was introduced to the array membrane and incubated overnight. Then, the relative amounts of 55 cytokines expression levels associated with angiogenesis were detected. Factors in red rectangles represent pro-angiogenic growth factors; and factors in blue rectangles represent anti-angiogenic growth factors (Figure 6). As the result, we found the pro-angiogenic proteins, Endothelin-1 and VEGF, and the anti-angiogenic proteins, Serpin E1 (PAI-1) were upregulated in both EGM-2 and HLFCM. In addition, the pro-angiogenic proteins, EGF, EG-VEGF, Persephin, bFGF, and the anti-angiogenic proteins, PF4 were upregulated in EGM-2 while downregulated in HLFCM. On the contrary, the pro-angiogenic proteins, IL-8, MMP-8, MMP-9, HB-EGF, Angiogenin, PDGF-AA, CXCL16, PIGF, and the anti-angiogenic proteins, Serpin F1 (PDEF), TIMP-1, TSP-1 were upregulated in HLFCM while downregulated in EGM-2.

**Figure 6.**
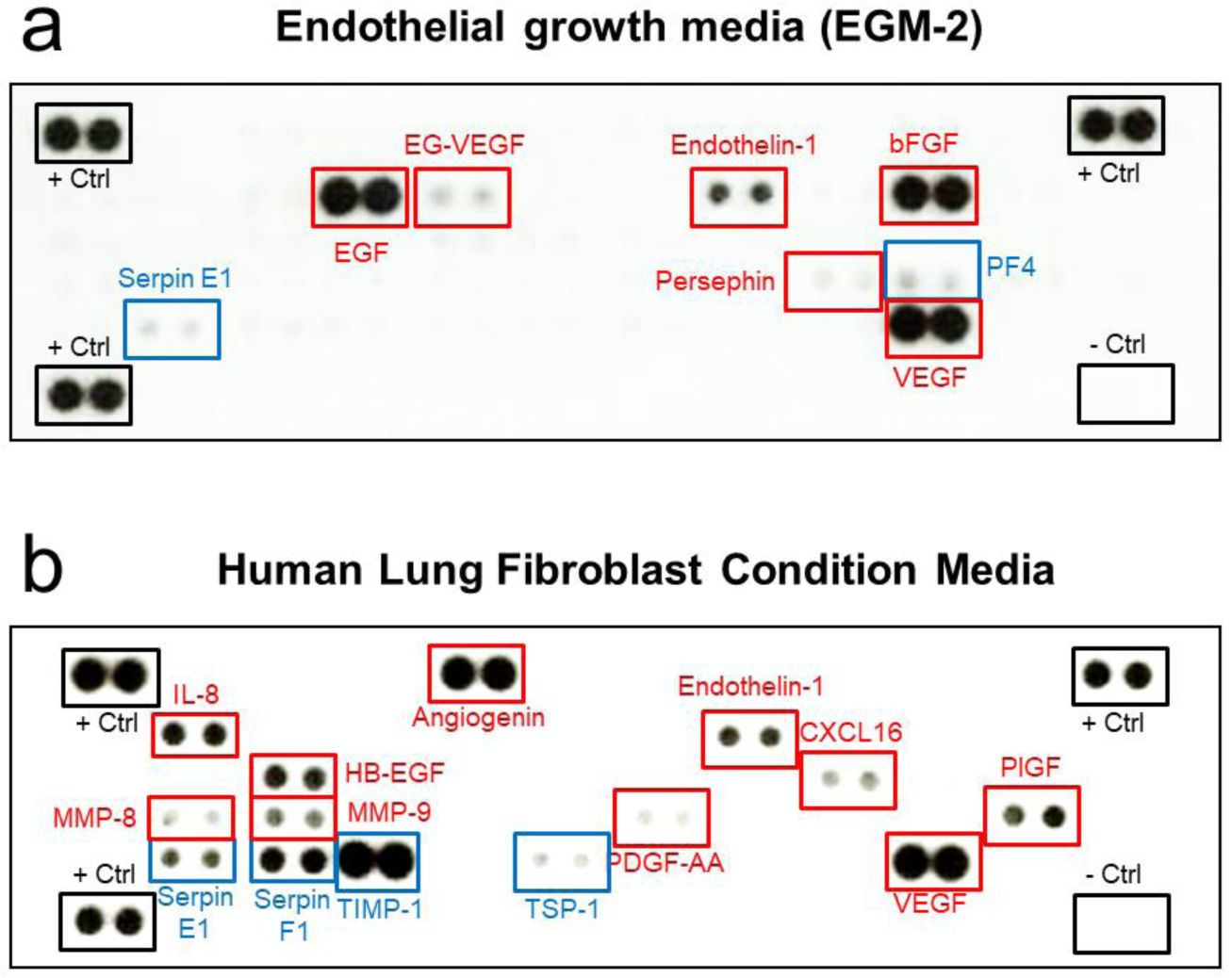
Reverse western assays. The reverse western assays with the human angiogenesis antibody arrays (R&D systems) detected the relative amounts of 55 cytokines in EGM-2 (a) and HLFCM (b). Representative images of the cytokine array membranes. Factors in red rectangles represent pro-angiogenic growth factors. Factors in blue rectangles represent anti-angiogenic growth factors.

## Discussion

We demonstrated the optimization process of engineered tumor spheroid-induced angiogenesis in a microfluidic organ-on-a-chip device to mimic vascularized breast tumor mass. We showed that among the ECMs used in the experiments, Matrigel allowed the migration of monoculture of MDA-MB-231 cells or mixed tumor spheroids of MDA-MB-231 cells and MSCs, but significantly disrupted the HUVEC channel by collapsing vascular lumen for unknown reasons (Figure 2). By contrast, collagen I allowed forming HUVEC lumens, however, the significant migrations of MDA-MB-231 cells in tumor spheroids were not observed (Figure 3). Interestingly, the fibrin ECM provided the both of forming HUVEC vascular lumens, angiogenesis, tumor invasion more actively than in any case where collagen I or Matrigel was used individually or mixed (Figure 4 and 5). The experimental cases of HLFs embedded in the ECM were showed more active angiogenetic behavior than no cell embedded ECM, and the comparison of the growth media showed that the HLF conditioned media included pro-angiogenic growth factors to encourage angiogenesis under the condition (Figure 6). These results strongly support the fact that HLFs in ECM have the ability to trigger angiogenesis. Consequently, it was found that a combination of HLFs laden fibrin-ECM, MDA-MB-231 cells and HUVECs mixed tumor spheroids was the most optimal cocktail to induce angiogenesis due to metastatic breast cancer with HUVEC engineered vascular structures among our trials. The reverse western array data revealing secreted factors in HLFs remain further investigation to identify potential new targets to treat tumor angiogenesis.

Feasibility of controlling the experimental complexity is crucial that is required in *in vitro* models of the complex biomimicking system. The findings of our study represent the potential of the tumor spheroids-vascular crosstalk in the field of cancer research because the 3D tumor spheroids/vascular on-a-chip models provide a more precise *in vivo* tumor microenvironment (TME) to study tumor growth and development as well as the interactions among tumors, adjacent blood vessels, and extracellular matrices. We believe that the application of these multiple approaches will support obtaining a systematic understanding of developing practical strategies for cancer treatments. Moreover, the use of the system in cancer research will be a potential for the development of more advanced personalized medicine development.

## Acknowledgement

T.J.K. and E.L. are supported by the Cornell University Start-up funds, and the Nancy and Peter Meinig Family Investigator funds. This work was performed in part at the Cornell NanoScale Facility (CNF), a member of the National Nanotechnology Coordinated Infrastructure (NNCI), which is supported by the National Science Foundation (Grant NNCI-1542081).

## Conflict of interest

The authors declare no financial or commercial conflict of interest.

## Availability of data and material

All data generated or analyzed during this study are included in this published article.

## Notes

### Competing Interest Statement

The authors have declared no competing interest.

## References

Antoni, D., Burckel, H., Josset, E., & Noel, G. (2015). Three-dimensional cell culture: a breakthrough in vivo. International Journal of Molecular Sciences, 16(3), 5517–5527.

Atiya, H., Frisbie, L., Pressimone, C., & Coffman, L. (2020). Mesenchymal Stem Cells in the Tumor Microenvironment. Adv Exp Med Biol, 1234, 31–42. doi: 10.1007/978-3-030-37184-5_3

Chung, M., Ahn, J., Son, K., Kim, S., & Jeon, N. L. (2017). Biomimetic Model of Tumor Microenvironment on Microfluidic Platform. Advanced Healthcare Materials, 6(15), 1700196. doi: 10.1002/adhm.201700196

D.H.T. Nguyen, E. Lee, S.A. Alimperti, R.J. Norgard, A. Wong, Lee, J. J. K., … Chen, C. S. (2019). A biomimetic pancreatic cancer on-chip reveals endothelial ablantion via ALK7 signaling. Science Advances, 5, eaav6789.

Fennema, E., Rivron, N., Rouwkema, J., van Blitterswijk, C., & de Boer, J. (2013). Spheroid culture as a tool for creating 3D complex tissues. Trends in Biotechnology, 31(2), 108–115. doi: https://doi.org/10.1016/j.tibtech.2012.12.003

Folkman, J. (1971). Tumor Angiogenesis: Therapeutic Implications. New England Journal of Medicine, 285(21), 1182–1186. doi: 10.1056/nejm197111182852108

Hassell, B. A., Goyal, G., Lee, E., Sontheimer-Phelps, A., Levy, O., Chen, C. S., & Ingber, D. E. (2017). Human Organ Chip Models Recapitulate Orthotopic Lung Cancer Growth, Therapeutic Responses, and Tumor Dormancy In Vitro. Cell Rep, 21(2), 508–516. doi: 10.1016/j.celrep.2017.09.043

Hirschhaeuser, F., Menne, H., Dittfeld, C., West, J., Mueller-Klieser, W., & Kunz-Schughart, L. A. (2010). Multicellular tumor spheroids: An underestimated tool is catching up again. Journal of Biotechnology, 148(1), 3–15. doi: https://doi.org/10.1016/j.jbiotec.2010.01.012

Kim, S., Chung, M., & Jeon, N. L. (2016). Three-dimensional biomimetic model to reconstitute sprouting lymphangiogenesis in vitro. Biomaterials, 78, 115–128. doi: 10.1016/j.biomaterials.2015.11.019

Ko, J., Ahn, J., Kim, S., Lee, Y., Lee, J., Park, D., & Jeon, N. L. (2019). Tumor spheroid-on-a-chip: a standardized microfluidic culture platform for investigating tumor angiogenesis. Lab on a Chip, 19(17), 2822–2833. doi: 10.1039/C9LC00140A

Kopanska, K. S., Bussonnier, M., Geraldo, S., Simon, A., Vignjevic, D., & Betz, T. (2015). Quantification of collagen contraction in three-dimensional cell culture. In Methods in cell biology (Vol. 125, pp. 353–372): Elsevier.

Kwak, T. J., Nam, Y. G., Najera, M. A., Lee, S. W., Strickler, J. R., & Chang, W.-J. (2016). Convex Grooves in Staggered Herringbone Mixer Improve Mixing Efficiency of Laminar Flow in Microchannel. PLOS ONE, 11(11), e0166068. doi: 10.1371/journal.pone.0166068

Laurens, N., Koolwijk, P., & De Maat, M. P. M. (2006). Fibrin structure and wound healing. Journal of Thrombosis and Haemostasis, 4(5), 932–939. doi: 10.1111/j.1538-7836.2006.01861.x

Lee, E., Fertig, E. J., Jin, K., Sukumar, S., Pandey, N. B., & Popel, A. S. (2014). Breast cancer cells condition lymphatic endothelial cells within pre-metastatic niches to promote metastasis. Nat Commun, 5, 4715. doi: 10.1038/ncomms5715

Lee, E., Pandey, N. B., & Popel, A. S. (2015). Crosstalk between cancer cells and blood endothelial and lymphatic endothelial cells in tumour and organ microenvironment. Expert Rev Mol Med, 17, e3. doi: 10.1017/erm.2015.2

Lee, E., Song, H. G., & Chen, C. S. (2016). Biomimetic on-a-chip platforms for studying cancer metastasis. Curr Opin Chem Eng, 11, 20–27. doi: 10.1016/j.coche.2015.12.001

Nashimoto, Y., Okada, R., Hanada, S., Arima, Y., Nishiyama, K., Miura, T., & Yokokawa, R. (2020). Vascularized cancer on a chip: The effect of perfusion on growth and drug delivery of tumor spheroid. Biomaterials, 229, 119547. doi: https://doi.org/10.1016/j.biomaterials.2019.119547

Polacheck, W. J., Kutys, M. L., Tefft, J. B., & Chen, C. S. (2019). Microfabricated blood vessels for modeling the vascular transport barrier. Nature Protocols, 14(5), 1425–1454. doi: 10.1038/s41596-019-0144-8

van Duinen, V., Zhu, D., Ramakers, C., van Zonneveld, A. J., Vulto, P., & Hankemeier, T. (2019). Perfused 3D angiogenic sprouting in a high-throughput in vitro platform. Angiogenesis, 22(1), 157–165. doi: 10.1007/s10456-018-9647-0

